# Ubiquitinome dynamics and regulation by DUB2 during *Leishmania* differentiation

**DOI:** 10.64898/2026.05.08.723729

**Authors:** Sergios Antoniou, Nathaniel G Jones, Vincent Geoghegan, Adam Dowle, Charlotte McNiven, Rachel Neish, Chris MacDonald, Anthony J Wilkinson, Jeremy C Mottram

**Affiliations:** York Biomedical Research Institute and Department of Biology, University of York, UK; Bioscience Technology Facility, Department of Biology, University of York, UK; York Structural Biology Laboratory, York Biomedical Research Institute and Department of Chemistry, University of York, UK

## Abstract

Leishmaniasis is caused by Leishmania parasites, which undergo cellular adaptation when transitioning from the insect stage (promastigote) to the mammalian stage (amastigote). While the ubiquitin-proteasome system (UPS) is vital for life cycle progression, global ubiquitination dynamics have remained unmapped. We established a quantitative ubiquitinomics workflow for *Leishmania mexicana*, identifying over 9,100 ubiquitination sites across 38% of the proteome, revealing thousands of stage-specific regulatory events. Promastigote-enriched sites associate with cell motility, while amastigote-enriched sites link to metabolism and glycosome organization. We identified extensive ubiquitination on UPS components, including the essential virulence factor deubiquitinase 2 (DUB2). Using inducible gene deletion and XL-BioID proximitomics, we identified 111 potential DUB2 substrates. High-confidence substrates include the E2 conjugating enzyme UBC2, which is required for differentiation, and SUMO, a critical regulator of ubiquitin crosstalk. The discovery of UBC2 as a substrate of DUB2 directly links ubiquitination with promastigote to amastigote differentiation. Our findings provide a comprehensive map of the *Leishmania* ubiquitinome and demonstrate that DUB2 acts as a pleiotropic regulator controlling post-translational modifications of essential proteins involved in life cycle progression.

## Introduction

Leishmaniasis is a spectrum of neglected tropical diseases caused by obligate intracellular, protozoan parasites of the *Leishmania* genus (*1*). This disease, which can be zoonotic, is transmitted between mammalian hosts by the bite of infected female sand flies (*Phlebotomus* in the Old World and *Lutzomyia* in the New World). The parasite’s complex digenetic cycle involves two main developmental forms: the motile, flagellated promastigote found in the sand fly gut and the compact, non-motile amastigote form that replicates within mammalian macrophages (*2*).

The morphological change between the promastigote and amastigote forms is accompanied by profound changes in gene expression (*3*, *4*) and protein abundance (*5*), allowing the parasite to adapt to different environments in the two hosts. For example, comparative proteomics has revealed stage-specific expression of over 1,100 proteins as *Leishmania mexicana* differentiates from procyclic promastigotes to intracellular amastigotes (*6*). This differentiation process relies heavily on mechanisms of protein turnover, including autophagy and lysosomal proteolysis (*7*, *8*). A key regulator of protein homeostasis in eukaryotes is the ubiquitin-proteasome system (UPS), in which ubiquitin is covalently attached to specific substrate proteins through an enzymatic cascade, targeting them for recognition and ATP-dependent degradation by the 26S proteasome (*9*, *10*). K48-linked polyubiquitination is the canonical signal, a chain of four or more ubiquitin molecules linked via lysine 48 that targets the protein for the 26S proteasome (*11*). K63-linked chains are unique because they typically do not signal for degradation; instead, they facilitate protein scaffolding in signaling pathways and DNA damage responses (*11*). Previous studies have established that core components of the UPS are essential for the survival and differentiation of *L. mexicana* (*12, 13*). However, how the parasite’s entire landscape of ubiquitinated proteins, the ubiquitinome, changes between the promastigote and amastigote stages remains unknown. Such a characterization would highlight the specific post-translational regulatory changes that drive life cycle progression.

The process of ubiquitination is tightly controlled by ubiquitination enzymes (E1, E2 and E3) and several classes of proteases, the deubiquitinating enzymes (DUBs), which cleave ubiquitin from target proteins. While humans possess ∼100 cysteine protease family DUBs, the *Leishmania* genome encodes about 20 (*12*, *14*). Among these, DUB2 is considered a key enzyme and potential drug target as it is essential for the survival of promastigotes and is critical for establishing infection in mouse models (*12*). Furthermore, DUB2 was previously reported to localise predominantly to the nucleus, where it interacts with proteins involved in transcription and chromatin dynamics (*12*). Despite its established essentiality and role in parasite survival, the specific molecular function and substrates of DUB2 are unknown.

To address this, we first performed a stage-specific ubiquitinome analysis of *L. mexicana* promastigotes and axenic amastigotes using a diGly mass-spectrometry-based, peptide-centric approach (*15*). This ubiquitinomics analysis revealed significant stage-specific regulation, identifying over 830 and 1,100 proteins selectively ubiquitinated in promastigotes and amastigotes, respectively. Intriguingly, the ubiquitination status of DUB2 itself was significantly enriched in the promastigote stage, suggesting a mechanism for its own post-translational regulation. Building on this observation, we next sought to define the essential function of DUB2 by identifying its substrates. We combined two mass-spectrometry approaches: (1) ubiquitinomics on *L. mexicana* promastigotes with inducible depletion of DUB2, and (2) characterization of the DUB2 proximitome using biotin-proximity labelling to complement its known interactome. By integrating these datasets, we identified six high-confidence DUB2 substrates, including SUMO and UBC2 (*13*, *16*). Collectively, our study provides the first comprehensive characterization of the *Leishmania* ubiquitinome and establishes a molecular link between DUB2, its novel substrates (SUMO and UBC2), and the essential processes of parasite survival and life cycle progression.

## Results

### Differences in the proteome and ubiquitinome of *Leishmania* promastigotes and amastigotes

We performed a label-free, quantitative mass-spectrometry analysis comparing the ubiquitinome of *L. mexicana* promastigotes and 4-day post-differentiation axenic amastigotes using a workflow based on ubiquitin DiGly enrichment (**Figure 1A**). In total we identified 9,727 and 9,149 ubiquitinated (=DiGly) peptides in promastigotes and amastigotes, respectively mapping to 3,323 and 2,994 proteins. Principal component analysis (PCA) demonstrated that the two life-cycle stages clustered distinctly, confirming significant differences between their ubiquitinomes (**Supplemental Figure 1A**). Limma statistical analysis, (adjusted *p*-value of 0.05 and log_2_ Fold Change of >1), revealed thousands of significantly regulated ubiquitination sites (**Figure 1B**). Promastigotes showed significant enrichment of 2,011 ubiquitination sites, across 1,109 proteins. Gene Ontology (GO) analysis indicated a strong enrichment for terms associated with microtubule-based functions, such as cell motility (**Supplemental Figure 1B**). Amastigotes showed enrichment of 1,544 ubiquitination sites, across 833 proteins, which were primarily associated with metabolic processes, including the metabolism of carbohydrates and lipids, and glycosome organization (**Supplemental Figure 1C**).

**Figure 1:**
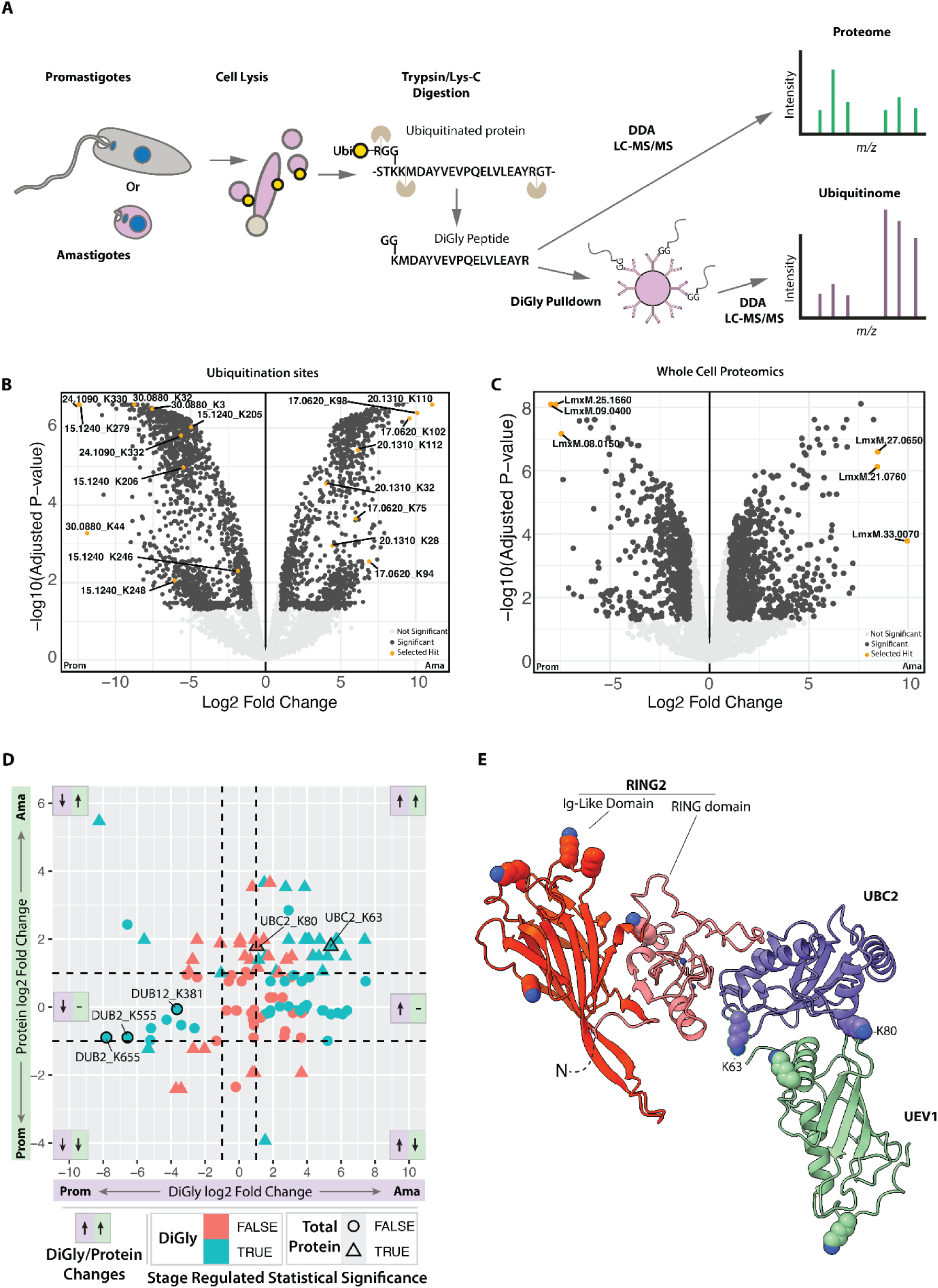
Significant ubiquitination sites and proteins enriched in *L. mexicana* promastigotes and amastigotes. **A.** Schematic of the ubiquitinomics/proteomics workflow **B.** Ubiquitination sites that are significantly different in promastigotes (Prom) and amastigotes (Ama), based on limma analysis of the ubiquitinomics dataset (adjusted *p*≤0.05; Log_2_FC>1). The top 3 changed ubiquitinated peptide sites are labelled with the protein ID and lysine residue number, as are the other ubiquitinated sites detected on the 3 proteins they are derived from. In this panel the LmxM prefix is omitted for clarity. **C.** Proteins enriched in promastigotes or amastigotes, based on limma analysis of the proteomics datasets. The top 3 differentially expressed proteins are labelled in each condition. **D**. Dot plot indicating the log2 fold-changes of 115 DiGly peptides (X-axis) against the levels of their parent protein in the whole-cell proteomics experiment (Y-axis). Stage regulated statistical significance is indicated by the teal colour and/or triangle symbol for the peptide and protein respectively, this is according to the pAdjust value from each individual analysis. Data are plotted for the 115 peptides identified in the DiGly dataset for which the protein was detected in the whole-cell proteomics. Positive log2 fold values indicate that there is a relative enrichment in the amastigote stage while negative values indicate enrichment in the promastigote stage. Dashed lines indicate a log2 fold change value of 1/-1. Panels indicate the relative fold change of the sectors, with up arrows indicating increase, a dash indicating stable, a down arrow indicating decrease, the purple sector for the DiGly site, and the green sector for the protein. A small selection of data points are labelled, corresponding to ubiquitinated peptides from DUB2, DUB12 and UBC2, with the residue number of the ubiquitinated lysine defined in the label. **E**. AlphaFold3 model of UBC2, UEV1 and RING2 depicted in ribbon format. Lysine residues that are detected as being ubiquitinated in one or more conditions are depicted as spheres. Zinc ions are modelled into RING2 using AlphaFold3. The disordered N-terminus of RING2 was omitted for visual clarity and is labelled N.

Stage regulated sites identified to be present on proteins in promastigotes, but not amastigotes, or vice versa numbered more than 2500. For promastigotes, examples include K32 on the hypothetical predicted multipass transmembrane protein (LmxM.24.1090), K279 on the nucleoside transporter 1 (LmxM.15.1240) and K44 on the amino acid permease 3 (LmxM.30.0880). Conversely, the most abundant ubiquitination sites on amastigote proteins were K110 on the small myristoylated protein 1 (LmxM.20.1310) and K98 and K102 in the Acyl CoA Binding Protein (LmxM.17.0620) (**Figure 1B, Supplemental Table 1**).

A parallel label-free quantitative proteomics analysis was performed on the same samples to assess changes in overall protein abundance. We identified a total of 3,776 and 3,801 proteins in the promastigote and amastigote samples, respectively. PCA again showed a clear segregation of the two life-stage clusters (**Supplemental Figure 1B**). Limma analysis identified a total of 1,369 proteins with significantly different abundance: 653 were upregulated in promastigotes and 716 in amastigotes (**Figure 1C**). Consistent with the ubiquitinome data, GO analysis indicated that promastigote-enriched proteins were associated with microtubule-based processes, while amastigote-enriched proteins were primarily involved in metabolic processes (**Supplemental Figure 1E &F**).

Top regulated proteins included the CLK1/KKT10 (LmxM.09.0400; 254-fold decrease in amastigotes), a hypothetical protein (LmxM.25.1660; 214-fold decrease), and the PIG-H component of the GPI-GlcNAc transferase complex (LmxM.08.0150; 176-fold decrease). In contrast, the top three most significantly enriched proteins in amastigotes were: ascorbate peroxidase (LmxM.33.0070; 996-fold increase), a hypothetical protein (LmxM.27.0650; 355-fold increase), and the proteasome regulatory non-ATPase subunit 5 (LmxM.21.0760; 350-fold increase) (**Figure 1C and Supplemental Table 1**).

The whole cell and DiGly proteomics datasets allowed us to explore the ubiquitination system of *Leishmania* for potential points of recursive regulation by ubiquitination (**Figure 1D**) exploring a cohort of 156 E1/E2/E3 enzymes, DUBs, Cullins, F-box proteins, SUMOylation enzymes and NEDD8 (**Supplemental Table 1**). Sixty-six proteins from this subset were found to contain one or more ubiquitination sites (totalling 151 distinct sites). Of these 151 sites, 79 were identified as differentially abundant between amastigotes and promastigotes, indicating a wealth of opportunities for recursive ubiquitin driven signalling events regulating stage-specific differentiation processes. In the whole cell proteomics, 82 proteins from the cohort were identified 44 of which were among the 66 detected in the DiGly proteomics analysis. These 44 proteins and their 115 sites of ubiquitination were examined in detail with respect to the fold changes in ubiquitination at each site and the parent protein abundance in the amastigote and promastigote stages (**Figure 1D**).

Firstly, instances where site-specific ubiquitination tracked with elevated total protein levels, potentially reflecting constitutive regulation, were examined. This totalled 22 ubiquitination sites on 8 proteins – UEV1, UBC2, DUB15, DUB18, PPDE5, HECT11, HECT12 and an unnamed RING protein (LmxM.12.0100). Intriguingly, HECT11 contains 14 ubiquitination sites with 9 that increase proportionally in the amastigote with increases in the protein levels, as well as a ubiquitination site (K4344) that decreases in abundance in the amastigote forms suggestive of differing specific regulatory events on the same protein. Also detected was the dual HECT ubiquitin ligase protein kinase TKUL (HECT12-LmxM.34.4000) with two ubiquitination sites that are significantly upregulated in the amastigote with the levels of total protein also significantly increased (*17*). The protein kinase and ubiquitin ligase activities of this protein are important for amastigote forms, and autophosphorylation of the protein is required for ubiquitin ligase activity. The discovery of these two sites raises the potential for further regulation of the protein’s function by ubiquitination.

Secondly, instances where a ubiquitinated peptide increased in the amastigote stage while the amount of total protein remained unchanged with respect to the promastigote stage (**Supplemental Table 1**) were evaluated. These represent ubiquitination events that could be correlated with stage-specific processes or potentially act as causal modifications in stage differentiation. Twenty-six sites were identified on 11 proteins, these being RING2, UBA1b, UBC1, UBC8, DUB9, DUB13 DUB17, ULP2, HECT9, HECT10 and an unnamed RING protein (LmxM.29.1230). Notably the E3 ligase RING2 was identified as having highly upregulated levels of ubiquitination at 4 sites (K164, K188, K247, K354). The K164 site was not detected at all in the promastigote sample group; the total protein levels of RING2 did not significantly change. RING2 is known to interact with the UBC2 and UEV1 E2 conjugating factors, which are necessary for stage differentiation and can catalyse the formation of K63 linked di-ubiquitin in vitro (*18*). This is highly suggestive of an *in vivo* role for a ubiquitination cascade proceeding through this complex that is essential for differentiation (*13*, *18*). Three of the four sites of ubiquitination occur on an Ig-like domain of RING2 predicted in AlphaFold models (**Figure 1E**), which suggests that ubiquitination at these sites may alter protein-protein interactions. The fourth ubiquitination site in RING2 is found in the unstructured C-terminal region. Interestingly, the E1 activating enzyme Uba1b was itself detected as having seven ubiquitin sites, all of which are upregulated in amastigotes but without the protein level changing. This factor represents an activation gateway for ubiquitin to enter subsequent conjugation and ligation steps and its ubiquitination may act as a throttle for downstream ubiquitin mediated signalling.

In addition to the observation that SUMO was ubiquitinated, we observed that the SUMO E1 (UBA2) and the SUMO processing factor ULP2 each contain a ubiquitination site that increased in abundance in the amastigote, while total protein levels remained stable – providing further options for layering and integrating the signalling events mediated by SUMO and ubiquitin.

Although the DiGly proteomics provides no information on the ubiquitin linkages directed after substrate modification, as K48-linked polyubiquitin would lead to proteasomal targeting and destruction of the target protein it is likely that we are detecting DiGly sites with other functional linkages - for example K63, which directs other regulatory functions. There was only a single instance of a site where ubiquitination increased as the corresponding total protein level decreased (LmxM.36.2930, an unnamed RING domain protein) (**Figure 1D**). Further experiments using proteasome inhibitors will be needed to properly explore the role of K48-linked ubiquitin and substrate degradation in the process of promastigote to amastigote differentiation.

Three ubiquitination sites were identified on two of the four essential *L. mexicana* DUBs. Two sites on DUB2 and one on DUB12 were significantly upregulated in promastigotes (**Figure 1D and Supplementary Figure 1E**), suggesting a stage-specific post-translational regulation mechanism for these key enzymes. While the overall abundance of the four essential DUBs did not change significantly, the observed stage-specific ubiquitination of DUB2 suggests regulation via post-translational modification, rather than protein degradation. Given the high enrichment of DUB2 ubiquitination in promastigotes and its known essentiality, we focused on DUB2 for subsequent functional analyses.

### Generation and validation of a DUB2 conditional knockout in *Leishmania* promastigotes

To investigate the essential function of DUB2 by identifying its substrates, we generated a DUB2::HA^Flox+/+^ promastigote cell line from a T7/Cas9 DiCre parental line. This system utilizes a modified rapamycin-DiCre-based inducible CRISPR-Cas9 approach (*19*, *20*) to replace both endogenous *DUB2* alleles with a *loxP*-flanked *DUB2::HA* construct. (**Figure 2A, Supplementary Figure 3A**). Successful expression of the DUB2::HA protein was achieved (**Supplementary Figure 3B**). The DUB2::HA^Flox+/+^ promastigote cell line showed a profound growth defect after incubation with rapamycin (RAP) compared to DMSO treatment (see example of clone A, **Figure 2B**). PCR-ba s ed genotyping and Western blot analysis confirmed the successful excision of the *DUB2*gene and the concomitant lo ss of DUB2 protein(**Figure 2C, D**). We concluded that four days of RAP treatment was optimal for downstream proteomics experiments, as it achieved significant DUB2 depletion with minimal cell death.

**Figure 2:**
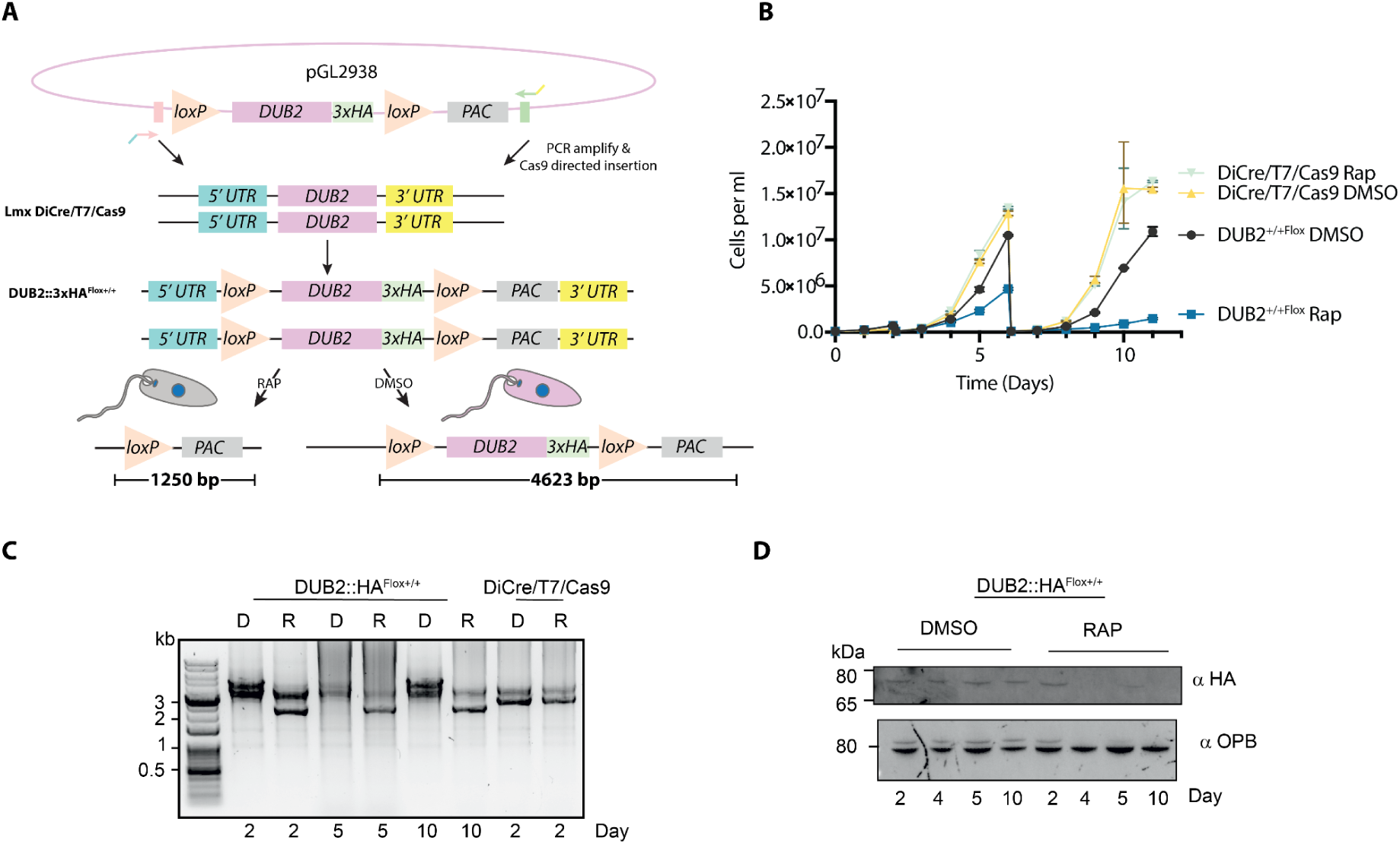
Generation of T7Cas9 DUB2::HA^Flox+/+^ *L. mexicana* promastigote cell line. **A.** Schematic of how the T7Cas9 DUB2::HA^Flox+/+^ promastigote cell line was generated and how rapamycin (RAP) induces excision of the *DUB2* gene. **B.** Growth curve of clone A (CLA) of T7Cas9 DUB2::HA^Flox+/+^ promastigotes following treatment with DMSO or rapamycin (RAP) (n=3). The parental T7Cas9 DiCre cell line (T7DiCre) was used as control (n=3). Cells were incubated for 11 days in the presence of either DMSO or RAP (300 nM), at a starting concentration of 1 x 10^5^ cells mL^-1^. At days 2 and 6, cells were re-seeded in fresh media, at a concentration of 1 x 10^5^ cells mL^-1^, and fresh DMSO or RAP (300 nM) was added. Data points represent means ± standard error of the mean (SEM). **C.** Genetic validation of DUB2 excision. The upper schematic illustrates the expected PCR amplicon sizes for rapamycin (RAP)- and DMSO-treated cells. The lower panel shows PCR products resolved on a 1% agarose gel using genomic DNA isolated from DUB2::HA^Flox+/+^ clone A and parental T7Cas9 DiCre promastigotes. Treatment conditions (D = DMSO; R = RAP) are indicated above each lane, and sampling time points (days) are indicated below the gel. **D.** Western blot to measure DUB2 protein (∼81 kDa) after DMSO and RAP treatment for 2, 4, 5 and 10 days. Primary α-HA rabbit antibody (1:5,000) and secondary donkey α-rabbit Dylight650 (1:5,000) were used. OPB was used as a loading control, detected by the primary α-LmjOPB antibody (1:20,000) combined with the secondary donkey α-sheep Dylight488 (1:5,000) antibody.

### Identifying DUB2 substrates via ubiquitome and proteome changes

Using the established inducible system (*20*) and experimental workflow shown in **Figure 3A**, we performed ubiquitinomics and proteomics on T7Cas9 DUB2::HA^Flox+/+^ promastigotes treated with DMSO or RAP for four days. PCA showed clear separation between the DMSO- and RAP-treated samples, confirming that DUB2 depletion drives the variance in the ubiquitinome and proteome profiles (**Supplementary Figure 2A, B**). For the ubiquitinomics analysis, we identified 7,757 ubiquitination sites, of which 6,605 were found in DMSO-treated samples, and 6,567 in RAP-treated samples, with ubiquitination sites on 2,173 and 2,159 proteins, respectively. Limma analysis revealed 174 significantly changed ubiquitination sites. Of these, 121 sites on 111 proteins were significantly increased in the absence of DUB2 (**Figure 3B; Supplementary Table 2**). This increase suggests that these proteins are likely DUB2 substrates that become hyper-ubiquitinated when the enzyme is absent. The top three increased ubiquitinated sites in the absence of DUB2 were K29 from the serine-threonine kinase receptor-associated protein (STRAP; LmxM.27.1250), K524 in the hypothetical protein (LmxM.34.3240) and K53 in the amino acid transporter 19 (AA19; LmxM.07.1160).

**Figure 3:**
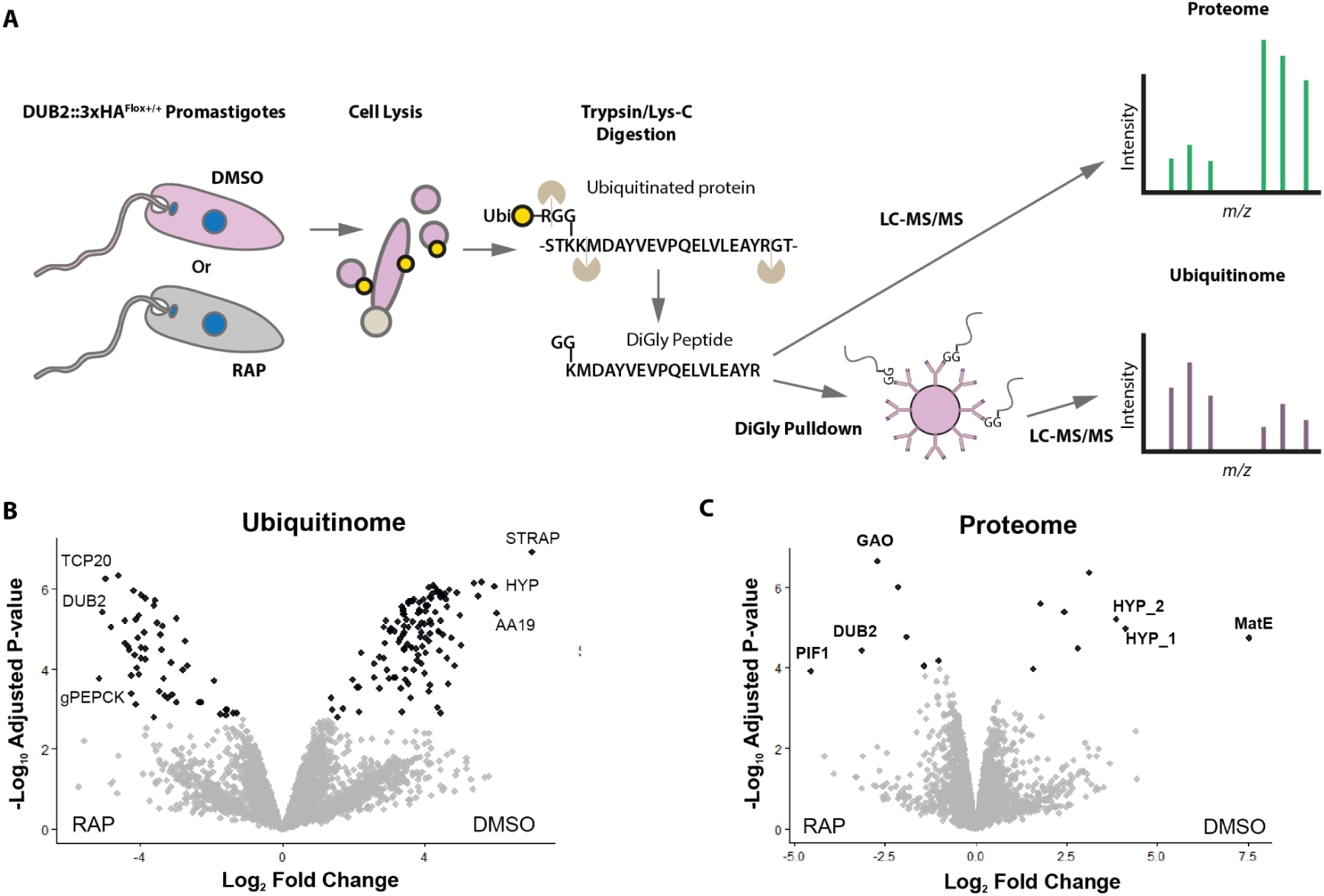
Ubiquitinomics and proteomics data in response to DUB2 depletion. **A.** Schematic of the ubiquitinomics and proteomics workflow, comparing RAP and DMSO-treated DUB2::HA^Flox+/+^ promastigotes. **B.** Limma analysis showed that ubiquitination on 121 sites was significantly increased upon RAP-induced DUB2 depletion (-DUB2), while 53 ubiquitination sites were significantly decreased (+DUB2; n=3). Significance corresponds to *p.adj*-values < 0.05 and Log_2_FC >1 represented by black dots. Ubiquitination sites that did not meet these criteria are depicted with grey dots. Abbreviations: TCP20, chaperonin TCP20 (LmxM.13.1660); gPEPCK, glycosomal phosphoenolpyruvate carboxykinase (LmxM.27.1805); STRAP, serine-threonine kinase receptor-associated protein (LmxM.27.1250), HYP, hypothetical protein (LmxM.34.3240); AA19, amino acid transporter 19 (LmxM.07.1160). **C.** Limma analysis revealed 8 proteins to be significantly increased and 7 to be significantly decreased in response to conditional DUB2 deletion (parameters as in **B**). Abbreviations: MatE (LmxM.34.3580); HYP_1 (LmxM.31.1380); HYP_2 (LmxM.26.1070); GAO (galactose oxidase LmxM.28.2150); PIF1, helicase-like protein (LmxM.21.1190).

A parallel proteomics experiment was conducted to determine how DUB2 depletion affects overall protein abundance. This analysis detected 5,748 proteins, of which 5,718 and 5,720 were enriched in DMSO- and RAP-treated samples, respectively. Limma analysis identified 15 proteins that were significantly changed, with 8 significantly depleted in the RAP-treated samples (**Figure 3C & Supplementary Figure 2D**). The most significantly depleted proteins were PIF1 helicase-like protein (LmxM.21.1190, 23-fold decrease), DUB2 itself (LmxM.08_29.2300, 9-fold decrease), and a hypothetical protein (LmxM.28.2150, 7-fold decrease). Notably, the significant decrease in DUB2 protein levels in RAP-treated compared to DMSO-treated samples validates the success of the conditional knockout system.

To rigorously select promising DUB2 substrates, we required that candidates have significantly increased occupancy of specific ubiquitination sites upon DUB2 depletion without a significant change in overall protein abundance. This criterion isolates changes driven by deubiquitination activity from changes driven by protein degradation. Of the 111 proteins with increased ubiquitination, 107 exhibited no significant change in overall protein abundance (**Supplementary Figure 2E**), making them promising candidates for DUB2 substrates.

### Characterisation of the DUB2 Proximitome

To complement the DUB2 ubiquitinomics and further inform substrate selection, we characterized the proximitome of DUB2 using XL-BioID (*21–23*). We generated a DUB2::miniTurbo::Myc cell line and used a nuclear-localized Myc::miniTurbo::CLK1 line as a compartment-specific control (**Figure 4A and Supplementary Figure 3A-C**). Biotinylated DUB2 and CLK1 were detected after addition of exogenous biotin (**Supplementary Figure 4A, B,** lane 6), however, biotinylation of these bait proteins was also observed in the absence of exogenous biotin (**Supplementary Figure 4A, B,** lane 8). We therefore refined the system using BioLock (*24*) to deplete biotin from the medium. Overnight BioLock treatment followed by 30 min of exposure to exogenous biotin effectively regulated miniTurbo activity (**Supplementary Figure 4A, B,** compare lanes 2 and 4) without significantly affecting cell viability for up to three days (**Supplementary Figure 4C,D**)

**Figure 4:**
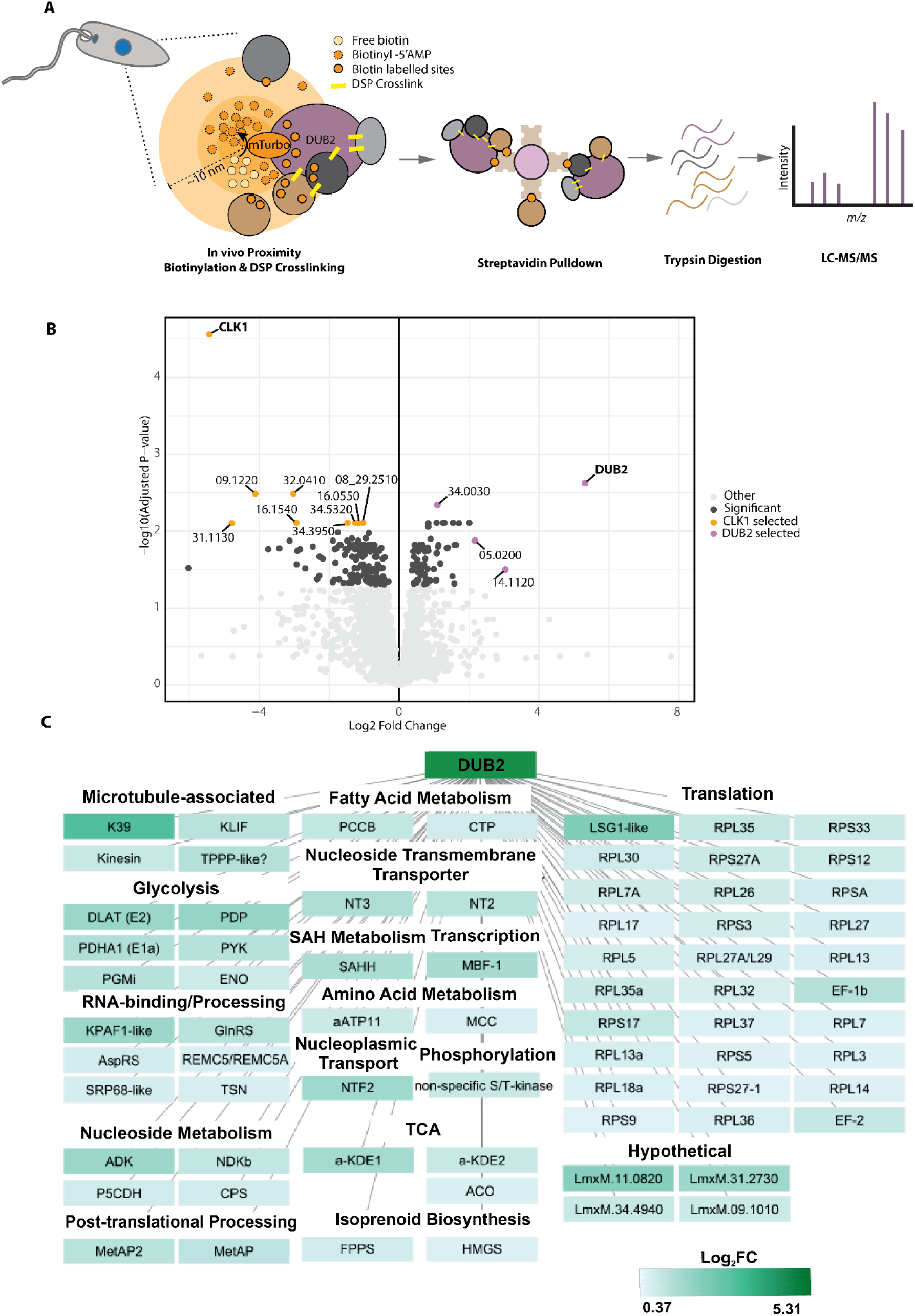
Investigation of the proximal interactome of DUB2. **A.** Schematic of proximity dependent biotinylation using miniTurbo-tagged DUB2 promastigotes is shown. An equivalent protocol was followed for the miniTurbo-tagged CLK1 promastigote cell line. **B.** Significantly enriched proteins in the DUB2 (73 hits; blue) and CLK1 (165 hits; red) datasets. Enrichment analysis was performed using the Limma package in Rstudio (*adjusted*.*p* <0.05; FDR= 5%; n=3). In grey are identified proteins that did not exceed the significance threshold. A selection of enriched proteins are labelled for each bait protein, coloured orange for CLK1 and purple for the DUB2 datasets, the LmxM. prefix is omitted for clarity. **C.** Proximal interactome of DUB2. Proteins were clustered based on their main biological process, in descending order from proteins with the highest Log2 Fold Change (FC; darker green) to proteins with the lowest Log_2_FC (light green). The extended version of the protein acronyms and their exact Log_2_FC value can be found in Supplementary Table 3.

Label-free tandem mass-spectrometry (MS/MS) analysis of the enriched biotinylated and cross-linked proteins revealed 73 proteins significantly enriched (*p.adj*-values < 0.05) in the DUB2 samples and 165 in the CLK1 control (**Figure 4B, Supplementary Table 3**). Confidence in the quality of our MS data was provided through PCA analysis (**Supplementary Figure 4E**), which revealed distinct clustering of DUB2::miniTurbo::Myc and Myc::miniTurbo::CLK1 samples. The top three enriched proximal proteins for DUB2 were DUB2 itself (40-fold), kinesin K39 (8.2-fold, LmxM.14.1120), and the 60S ribosome-binding GTPase (LSG1 LmxM.05.0200) (4.5-fold).

Gene ontology (GO) enrichment analysis showed that the proximitome is associated with 17 different biological processes, with protein translation having the most hits (41%), followed by RNA binding/processing (8%), cytoskeletal-associated proteins (7%) and glycolysis (7%; **Figure 4C**). We compared the XL-BioID proximal interactome with DUB2 co-immune-precipitation data previously generated by Damianou *et al.* (2020). Six of these DUB2-interacting proteins were also present in the proximal interactome, these being DUB2 itself (**Figure 4C**) and five proteins from the large (60S) ribosomal subunit L13a (RPL13a; LmxM.15.0200), L14 (RPL14; LmxM.22.1520), L7 (RPL7; LmxM.26.0170), L30 (RPL30; LmxM.34.0240) and L18a (RPL18a; LmxM.34.0600).

### Selection of high confidence substrates and validation

Promising DUB2 substrates were selected by integrating three datasets. From this study, firstly, the DUB2 proximity-labelled proteins and secondly proteins with significantly increased ubiquitination upon DUB2 depletion. The third dataset was the DUB2 interacting proteins from Damianou et al., 2020 (*12*). This identified six proteins that were present in at least one interactomics dataset and also showed significantly increased ubiquitination upon DUB2 depletion (**Figure 5A; Supplementary Table 4**); SUMO (LmxM.08.0470), UBC2 (LmxM.04.0680), iPGMA (LmxM.36.6650), CPSF73 (LmxM.33.3430), RPSA (LmxM.36.5010), and GDI (LmxM.08_29.2160).

**Figure 5:**
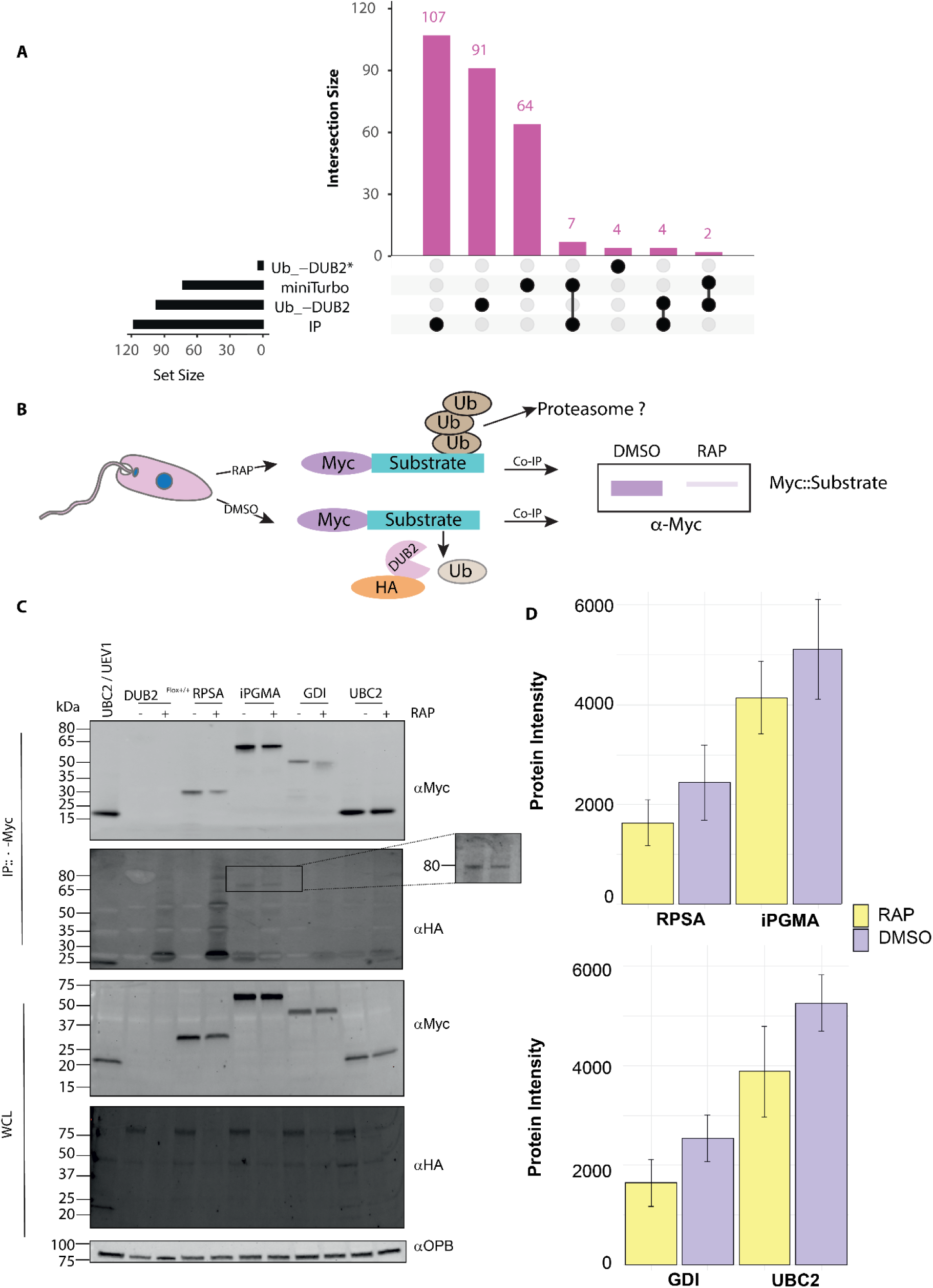
Selection and validation of promising DUB2 substrates: **A.** Overlap of significantly enriched proteins derived from the following datasets: the XL-miniTurboID-derived proximitome (miniTurbo); significantly increased ubiquitination sites in response to rapamycin-induced DUB2 depletion occurring on proteins whose overall abundance remained stable (Ub-DUB2); significantly increased ubiquitination sites upon DUB2 depletion, where total protein data was unavailable but other sites on the same protein remained unchanged, suggesting site-specific regulation (Ub-DUB2*); and the DUB2-co-immunoprecipitation dataset generated by Damianou et al. (2020; IP). Black dots indicate the combinations of datasets being compared, while the number of each of these proteins shared in that combination is plotted as a column chart (intersection size) with the number of shared proteins written on top of each column. The number of proteins found in each of the datasets being compared is presented as a bar chart (left-hand side). **B.** Schematic of the experimental procedure testing whether the Myc-tagged substrate protein levels of the investigated hits change in response to DUB2 depletion (RAP: rapamycin; Co-IP: co-immunoprecipitation). Rapamycin-induced deletion of DUB2 is expected to lead to increased levels of the substrate if the ubiquitin linkages removed by DUB2 are those that target proteins for proteasomal degradation (e.g. K48 polyubiquitin). **C.** Western blot analysis revealed the enrichment of MYC-tagged candidate substrates, in response to either RAP or DMSO treatment (IP: immunoprecipitated). Re-probing of the same membrane using anti-HA revealed no co-immunoprecipitation of DUB2, except for the case of iPGMA under higher image exposure. Pre-immunoprecipitated samples (WCL: whole cell lysate) validated the presence of Myc-tagged substrates, while re-probing of the same membrane with an anti-HA showed that rapamycin-induced DUB2 depletion was successful. Oligopeptidase B protein was used as loading control. The parental cell line DUB2::HA^Flox+/+^ was used as a DUB2-depletion control, and a UEV1::HA/UBC2::Myc cell line was used as a co-immunoprecipitation control. D. t-test analysis of band intensities, normalised to the loading control OPB from the input, revealed no significant changes in the enriched DUB2 substrate proteins following DUB2 depletion (p > 0.05; n = 4).. Abbreviations: Rab GDP dissociation inhibitor (GDI); 2,3-bisphosphoglycerate-independent phosphoglycerate mutase (iPGMA); ribosomal protein SA (RPSA); E2-conjugating enzyme (UBC2).

To further characterise these candidate substrates, we endogenously Myc-tagged each of the six proteins in the DUB2::HA^Flox+/+^ parental line and analysed their protein levels after DUB2 depletion (**Figure 5B**). Western blot analysis showed that none of the six candidates had a statistically significant change in overall protein abundance upon DUB2 depletion (**Figure 5C, D**), consistent with the proteomics data. This suggests that DUB2 deubiquitination of these substrates does not rescue them from proteasomal degradation, implying that for these substrates ubiquitination is associated with control of other cellular functions. An alternative explanation is that the proportion of ubiquitinated substrate in the total pool is too small to be measured by the Western blot assay.

## Discussion

### A comprehensive *Leishmania* ubiquitinome reveals stage-specific regulation

The ubiquitin-proteasome system (UPS) is a central element in the differentiation of the digenetic *Leishmania* parasite (*8*, *12*, *13*). However, the specific regulatory changes taking place at the post-translational level have remained undefined. In this study, we established the first quantitative ubiquitinomics pipeline in *L. mexicana* under native conditions, bypassing the need for proteasomal inhibitors like MG132, allowing profiling of the parasite ubiquitinome, and the identification of stage-specific changes between promastigotes and amastigotes. Our comparative analysis identified 9,727 and 9,149 ubiquitination sites in promastigotes and amastigotes, on 3,323 and 2,994 proteins respectively, representing nearly 38% of the predicted *L. mexicana* encoded genome. The finding that the majority of ubiquitination sites are present in both life stages suggests their involvement in fundamental, constitutively active molecular processes, similar to observations on phosphorylation (*25*). Nevertheless, we identified 2,011 and 1,544 significantly enriched ubiquitination sites in promastigotes and amastigotes, respectively. These stage-specific patterns reveal global post-translational regulation. Promastigote-enriched sites were primarily associated with microtubule-based functions (e.g., cell motility), consistent with the flagellated, motile form, while amastigote-enriched sites clustered in metabolic processes (e.g., lipid and carbohydrate metabolism) and peroxisome/glycosome organization, reflecting the parasite’s metabolic adaptation inside the mammalian host macrophage (*8*, *26–29*).

Intriguingly, our stage-specific ubiquitinome data revealed that the essential deubiquitinases DUB2 and DUB12 are themselves regulated by ubiquitination, with multiple sites significantly upregulated in promastigotes despite stable protein levels. This suggests that ubiquitination modulates DUB activity or localization in a stage-specific manner, rather than targeting them for degradation. As DUB2 is a key virulence factor and its ubiquitination status is highest in the promastigote stage, we focused on defining its molecular function.

### DUB2 depletion defines a pleiotropic regulatory role

To address the mechanism behind DUB2’s essentiality, we combined a DUB2 inducible knockout system (DUB2^Flox+/+^) with a rigorous, multi-faceted mass-spectrometry approach based on the principle that the absence of DUB2 will lead to the hyper-ubiquitination of its substrates (*30*). Depleting DUB2 in promastigotes for four days led to the significant hyper-ubiquitination of 121 sites across 111 proteins, strongly suggesting they are direct or indirect DUB2 substrates. Specific, highly regulated targets, such as the serine-threonine kinase receptor-associated protein (STRAP) and the amino acid transporter 19 (AAT19), are novel ubiquitination targets in *Leishmania*. Given the known roles of their eukaryotic homologues in apoptosis and nutrient signalling (*31*, *32*), it will be important to explore how DUB2 activity dynamically regulates these functions.

To complement the substrate predictions, we mapped the DUB2 proximitome using XL-BioID. We optimized this proximity labelling technique by introducing a BioLock pre-incubation step to control high background biotinylation—a necessary optimization given the parasite’s high-affinity biotin uptake systems and the constitutive activity of the miniTurbo enzyme (*33*). This approach identified 73 proximal partners, with kinesin K39 and ribosomal LSG1 as top hits. The association with Kinesin K39 suggests a role for DUB2 in cell-cycle-dependent trafficking and cytokinesis, which aligns with conserved functions observed for human DUB2 homologues USP13 and USP5 (*34*, *35*). Furthermore, the enrichment of five ribosomal proteins strongly supports a primary role for DUB2 in regulating translation and ribosome-associated processes, consistent with other kinetoplastid DUBs (Brannigan et al., 2025; Meyer et al., 2020). DUB association with ribosomes is well established, as USP10 was found to act in ribosome-associated quality control, ensuring the rescue of functional ribosomal proteins from proteasomal degradation (Jung et al., 2017; Meyer et al., 2020). The role of ubiquitination could mirror that of differential pseudouridination of ribosomal RNAs in driving stage-specific ribosomal function and warrants further investigation (*36*, *37*). Furthermore, DUB16 from *L. donovani* has been shown to cleave the ubiquitin-ribosomal L40 precursor protein in vitro (*38*). The identification of partners involved in 17 distinct biological processes reinforces the concept that DUB2 has evolved pleiotropic functions.

### Defining DUB2 substrates

By integrating the ubiquitinome, proteome, and interactome/proximitome datasets, we identified a high-confidence list of six DUB2 substrates. These candidates were selected using the criteria of being proximal to, or interacting with, DUB2, as well as exhibiting increased ubiquitination upon its depletion, while their overall protein levels remained stable. The latter point is important as it suggests DUB2’s role is primarily regulatory, mediating the cleavage of non-proteasome-directed ubiquitin, as opposed to stabilizing, rescuing proteins from degradation by cleaving proteasome-directing ubiquitin chains. The six candidates are Rab GDI, iPGMA, UBC2, CPSF73, RPSA, and SUMO. 2,3-bisphosphoglycerate-independent phosphoglycerate mutase (iPGMA) with its role in glycolysis/gluconeogenesis and Rab GDP dissociation inhibitor (GDI) with its role in trafficking are novel ubiquitination targets in *Leishmania*, positioning DUB2 as a regulator of core energy and trafficking pathways (*39*, *40*). Rab GDI, an evolutionarily conserved regulator of Rab proteins, controls intracellular trafficking by recycling inactive GDP-bound Rabs and delivering active GTP-bound Rabs to membranes. *Leishmania* possesses a single isoform of this enzyme in contrast to the three found in mammals (*39*, *41*). CPSF73 is an mRNA cleavage and polyadenylation factor, and it is ubiquitinated in yeast and humans by UBE3D to target it for degradation (*42–44*). Our data reveal multiple sites of CPFS73 ubiquitination in *Leishmania*, some of which may be involved in proteasome-directed degradation. Indeed, CPSF73 is thought to be the target of the anti-trypanosomal benzoxaboroles (*45*) and acoziborole (*46*). Targeted protein degradation of the CPSF complex, including CPSF73, by benzoxaboroles occurs through sumoylation and proteasome-dependent degradation (*47*). The identification of SUMO as a DUB2 substrate, with two potential ubiquitination sites, suggests DUB2 may function as a SUMO-targeted ubiquitin protease (STUP) (*48*). This places DUB2 at the nexus of the critical SUMOylation–ubiquitination crosstalk required for parasite survival (*16*, *49*). RPSA, a multifunctional ribosomal and signalling protein, has been linked to ubiquitination-dependent stability in mammals and is associated with RNA-binding proteins and telomerase components in kinetoplastids (*50–52*). This suggests that DUB2 is also involved in translation and chromatin regulation.

The experimental validation showing no change in protein abundance for these six proteins upon DUB2 depletion strongly supports their designation as true substrates, whose function is regulated by ubiquitination rather than being marked for degradation by it. This is further supported by the lack of stable co-immunoprecipitation of DUB2 with any of the candidates, suggesting transient or weak interactions characteristic of substrates, which are best captured by the XL-BioID proximity approach rather than traditional IP (*53*). Many deubiquitinases are known to be components of stable complexes in which their activity is regulated. For instance, the yeast DUBs Ubp8 and Rpn11 are components of the histone-modifying complex SAGA and the 26S proteasome, respectively, and association with their interacting partners (e.g. Sgf11 and Rpt1, respectively) induces their activation (*54*, *55*)). This does not appear to be the case for DUB2. The evidence supports the concept that DUB2 functions autonomously within multiple compartments of the cell, and the examined interacting proteins are more likely to serve as substrates of DUB2, due to their transient and weak interaction with it.

### Regulation of UBC2 activity during differentiation

UBC2 is an E2 conjugating enzyme essential for promastigote to amastigote differentiation (*13*). Consequently, DUB2’s activity on UBC2 provides a direct regulatory link between a deubiquitinating enzyme and the control of developmental transitions. *In vitro* in the presence of the E1 enzyme UbA1a, and ATP, the C-terminal carboxylate of ubiquitin becomes covalently linked to the active site Cys85 of UBC2. This donor ubiquitin (Ub^D^) complex is competent for RING E3 ligase catalysed transfer reactions leading to formation of either free, or target protein-linked, polyubiquitin chains (*13*). These reactions occur in the absence of the ubiquitin-conjugating enzyme variant, UEV1. In its presence, a stable UBC2-UEV1 complex is formed. This allows UBC2 to become similarly ubiquitinated on Cys85, however UEV1 controls the binding of the acceptor species, restricting it to a second molecule of ubiquitin (Ub^A^), and directing its lysine 63 side chain towards the activated donor ubiquitin resulting in formation of K63-linked diubiquitin (*13*).

The ubiquitinomics data reveal ubiquitination of UBC2 on K63 (not to be confused with K63 of Ub) and K80. The side chains of K63 and K80 flank the active cysteine residue (C85) and are oriented towards UEV1 in the crystal structure of the UBC2-UEV1 complex (**Figure 6A**, Burge 2020). It is therefore plausible that ubiquitination of UBC2 at one or both of these sites inhibits its E2 activity or changes its substrate and/or linkage specificity. The mode of ubiquitin binding in *L. mexicana* UBC2-UEV1 may be inferred from comparison with the structure of the homologous UBC2-Mms2 complex from *S. cerevisiae* crystallised with bound ubiquitin (*56*). As shown in **Figure 6B & C**, K63, which corresponds to E65 in the yeast homologue, projects towards Ub^D^ such that regulatory ubiquitination of this site might block thioester bond formation at C85 and inhibit the E2 activity. However, E2∼Ub^D^ complexes are known to have open and dynamic structures as a result of the extended, flexible C-terminal tail on the ubiquitin (*57*). RING E3 ligases serve to stabilise closed E2∼Ub^D^ conformations which are allosterically activated for Ub transfer (*58*). This activated conformation of UBC2∼Ub^D^ may be inferred from the structure of RNF4-RING E3 complexed with human Ubc13 and UbeV2 (**Figure 6D**, (*59*)). In this closed state, Ub^D^ is distally located so that the effects of regulatory ubiquitination at K63 on UBC2 would be less severe. It seems more likely therefore that regulatory ubiquitination of UBC2 alters access of acceptor species to the reactive centre. Acceptor ubiquitin packs onto the surface of UEV1, so that its K63 side chain is directed towards the active site thioester linked C85 (**Figure 6C**). The data presented here support the suggestion that ubiquitin modification of K63 of *L. mexicana* UBC2 leads to occlusion of the Ub^A^ binding surface of UEV1 allowing cognate E3 ligases, including RING2, to determine the identity and binding orientation of alternative acceptor species leading to altered cellular outcomes.

**Figure 6.**
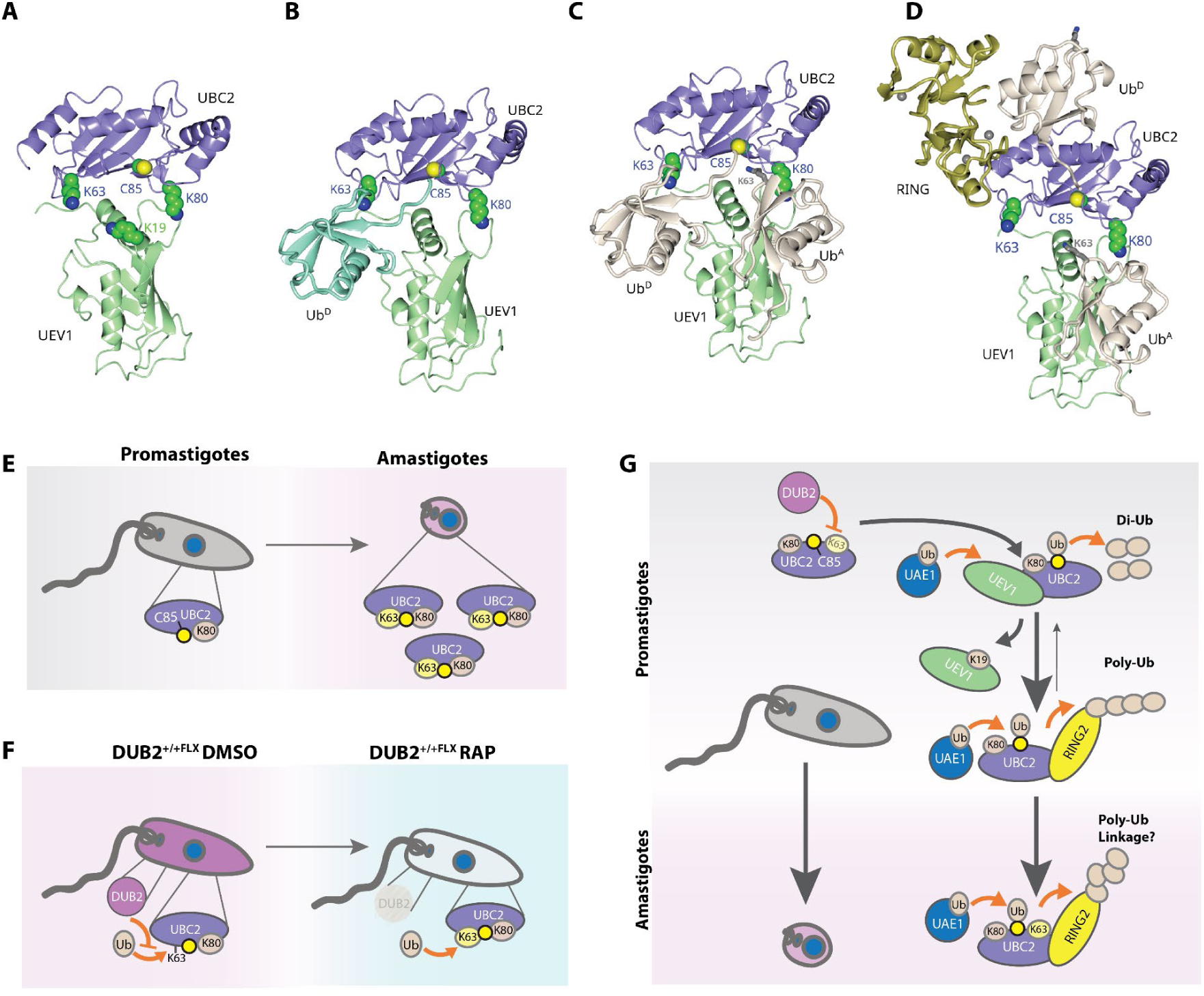
Conceptual model of UBC2 regulation by ubiquitin and DUB2. **A.** Crystal structure of the UBC2-UEV1 complex from *L. mexicana* (6zm1, (*13*)) with side chains of residues discussed in the text shown in sphere format. **B** & **C** Superposition onto this structure of donor (Ub^D^) alone (**B**), or donor and acceptor (Ub^A^) (**C**) ubiquitins from the structure of the yeast Ubc13-Ub-Mms1 complex (2gmi, (*56*)). The binding of Ub^A^ to UEV1 directs the side chain of K63 (shown in grey cylinder format) towards the active site Cys85 to which Ub^D^ is linked. In **D**, the RING E3 ligase domain of RNF4 and ubiquitin moieties (Ub^D^ and Ub^A^) from the Ubc13∼Ub-UbeV2-RNF4-RING structure (5ait, (*59*)) are displayed with UBC2-UEV1 following superpositioning based on the human E2 entities Ubc13 and UbeV2. **E.** Summary of UBC2 protein levels increasing during differentiation with concomitant stable levels of UBC2 K80 ubiquitination and increasing levels of UBC2 K63 ubiquitination. The ubiquitin attached to K63 of UBC2 is coloured light yellow to indicate it is stage-dependent. **F.** In promastigotes, when DUB2 is deleted, UBC2 levels remain stable but the levels of ubiquitination at the K63 residue are found to increase, with K80 ubiquitination also remaining stable (E and F show that K63 modification is stage-specific and DUB2-responsive). **G.** Conceptual model integrating Panels A-F to propose a hypothesis where DUB2 removes a constitutively added ubiquitin at the K63 site on UBC2 in promastigotes., Upon differentiation, this changes allowing the UBC2 K63 site to remain ubiquitinated. The interplay of ubiquitination of UEV1 K19, UBC2 K63 and K80 drives changes in the proportion of UBC2 pool partitioning into the UBC2:UEV1 and UBC2:RING complexes (indicated here by RING2). These may serve to modulate the cellular pools of di-ubiquitin and also the linkage specific ubiquitination conducted by downstream UBC2:RING ligase complexes, driving the ubiquitin-dependent processes that are essential for amastigogenesis. Orange arrows denote the flow of ubiquitin, grey arrows indicate change in the composition of protein complexes (or in the case of the parasites, life stage).

A third site of ubiquitination, K19 on UEV1, is located close to the UBC2 interface and to the sites of Ub^D^ and Ub^A^ binding (**Figure 6A**). UBC2 protein abundance and K80 ubiquitination increase concomitantly in amastigotes (**Figure 6E**), suggesting constitutive modification, while the K63 site displays stage-specific regulation. K63 ubiquitination is undetectable in promastigotes but is prevalent in the amastigote stage, and together with K63 being present in the absence of DUB2, these data provide evidence that DUB2 removes this modification (**Figure 6F**). Interestingly, while K80 is conserved across kinetoplastids, K63 is unique to *Leishmania*, suggesting a genus-specific regulatory function. Similar to K80 in UBC2, ubiquitination at K19 on UEV is not stage regulated suggesting a constitutive regulatory role.

Overall, the 2 ubiquitination sites at UBC2’s interface with UEV1; K63 and K80 (**Figure 1E & Figure 6E**) provide a potential mechanism by which the ratio of UBC2:UEV1 and UBC2:RING complexes can be varied dynamically (for example by DUB2 **Figure 6F)**, and thus determine the outcome of the downstream ubiquitin ligase activities (**Figure 6**). More detailed characterisation of the occupancy of ubiquitination at these sites could inform on the stoichiometry of these different cellular pools of UBC2:UEV1 and their relative contribution to diubiquitin or polyubiquitin formation.

In conclusion, this study provides the first comprehensive ubiquitinomics database for *L. mexicana*, identifying extensive stage-specific regulation and, critically, demonstrating that the essential DUB2 is itself post-translationally regulated. By leveraging a targeted, multi-omics approach, we propose six high-confidence substrates, including the essential proteins UBC2 and SUMO, that position DUB2 as a pleiotropic regulator of differentiation, metabolism, and gene expression.

## Methods

### Cell culture

*L. mexicana* (MNYC/BZ/62/M379) promastigote cell lines were grown at 25°C, in HOMEM medium (modified Eagle’s medium; Gibco) supplemented with 10% v/v heat-inactivated fetal bovine serum (FBS; Gibco) and 1% v/v penicillin-streptomycin solution (Sigma-Aldrich). Cells were kept at log-phase, unless otherwise stated. Drugs used for selection were added at the following concentrations (Beneke et al., 2017): 10 μg mL^-1^ blasticidin (InvivoGen), 15 μg mL^-1^ G418 (InvivoGen), 50 μg mL^-1-^hygromycin (InvivoGen), 50 μg mL^-1^ nourseothricin (Jena Bioscience) and 40 μg mL^-1^ puromycin (InvivoGen). For promastigote to amastigote differentiation, promastigotes were incubated for 3-4 days in stationary phase before harvesting at 1,000 x *g* for 10 min. Pelleted cells were resuspended in amastigote medium (pH 5.5) containing Schneider’s Drosophila medium (Gibco), 20% FBS (Gibco) and 15 μg mL^-1^ Hemin (Sigma). Cells were incubated at 33°C with 5% CO_2_ in either 6-well plates or vented flasks and maintained at 1 x 10^7^ cells mL^-1^.

### Endogenous tagging

For endogenous tagging of *L. mexicana* promastigotes, we used the CRISPR-Cas9-based approach developed by Beneke et al. (2017) using *L. mexicana* promastigotes expressing Cas9 and T7RNAP (T7/Cas9). Two sets of primers were designed per gene-of interest (GOI) through the online resource LeishGEdit (http://www.leishgedit.net): the first for generating the single guide RNA (sgRNA) template, and the second to amplify the DNA repair cassettes. The sgDNA scaffold was generated by PCR containing in 20 μl: 2 μM forward primer, 2 μM reverse OL6137 primer, 0.2 mM dNTPs, 1U Q5 DNA Polymerase (NEB), 1 X Q5 reaction buffer (NEB), and deionised H_2_O. The following PCR protocol was employed: 98°C for 30 sec followed by 35 cycles of 98°C for 10 sec, 60°C for 30 sec, and 72°C for 15 sec. A final extension at 72°C for 10 min was carried out. In the case of DNA donor cassette, the following PCR reaction was performed in a final volume of 40 μl: 2 μM forward primer, 2 μM reverse primer, 0.2 mM dNTPs, 1U Q5 DNA Polymerase (NEB), 1 X Q5 reaction buffer (NEB), 30 ng of a pPLOTv1 plasmid and deionised H_2_O. The PCR conditions for this PCR reaction were: 94°C for 5 min, 45 cycles of 94°C for 30 s, 65°C for 30 sec, and 72°C for 2 min 15s, followed by a final extension at 72°C for 7 min. PCR amplicons were validated via gel electrophoresis; sgRNA and DNA donor cassettes were mixed and then purified and concentrated via ethanol precipitation. Finally, they were transfected in the parental cell line, as described by Geoghegan *et al*. (2022). Primers and plasmids used are listed in Table S5.

### DiCre T7 Cas9 cell line and rapamycin-Induced knockout

To generate the inducible DiCre *DUB2* cell line, the CRISPR-Cas9-based endogenous tagging approach was utilized to replace the endogenous *DUB2* with a loxP-flanked sequence encoding *DUB2* C-terminally fused to a 3xHA tag cassette upstream of a puromycin resistance gene (Figure 2A). The DiCre T7/Cas9 promastigote cell line was used as the parental line (*22*). The donor cassette was generated via Gibson assembly. Briefly, the *L. mexicana* DUB2 gene was amplified via PCR using primers designed by NEBuilder. The pGL2938 plasmid was linearized with *Eco*RI and ligated with the *DUB2* amplicon for 1 h at 50°C. Sequentially, transformation of chemically competent *E. coli* cells was performed, plasmid was extracted via miniprep (Qiagen), and validated via whole-plasmid sequencing (Plasmidsaurus).

Subsequently, the loxP-flanked DUB2 C-terminally HA tagged cassette containing a puromycin resistance expressing gene was amplified from the plasmid, using primers containing homology arms that recombine at the 5’ and 3’ UTR of the endogenous *DUB2* and transfected within the parental cell line. Successful gene replacement was achieved via the use of two sgRNAs, which targeted the 5’ and 3’ UTR of endogenous *DUB2*, and cloned, followed by diagnostic PCR to ensure double allele replacement. All primers and plasmids are in **Supplementary Table 5**.

To excise *DUB2* from the homozygous mutants log-phase cells were seeded at 1-5 x 10^5^ cells mL ^-1^ into HOMEM medium and incubated for 2 d in the presence of 300 nM of rapamycin. Cells were then re-seeded at the desired cell density and incubated for 2-3 d in fresh HOMEM media containing 300 nM rapamycin. Cells were harvested at 1,000 x *g* for 10 min and prepared accordingly for downstream analyses. To determine the growth rate of the transgenic cell lines, cells were seeded at 1 x 10^5^ cells mL ^-1^ in HOMEM media and cell counts were made every 24 h for 5 d. Growth curves were plotted and analysed using Rstudio.

### Western blotting and protein quantification

Samples were mixed with 1 x NuPAGE LDS Sample Buffer containing 10 mM dithiothreitol (DTT) as a reducing agent, heated for 10 min at 70°C and loaded into NuPAGE™ 4-12% Bis-Tris Protein Gels (Invitrogen). Electrophoresis was carried out at 150 V for 1-2 h in 1 x NuPAGE™ MOPS SDS Running Buffer (Invitrogen). Then, gels were incubated in 10 mL of 20% ethanol for 10 min. Blotting was performed using PVDF iBlot™ 2 Transfer Stacks (Invitrogen) and the dry blotting system iBlot™ 2 (Thermo Fisher Scientific). Membranes were then blocked with either 5% skimmed milk in Tris-buffered saline containing 0.1% Tween 20 detergent (TBST) or 5% BSA (Merck) for 1 h. Probing with primary antibodies diluted in either 5 % milk TBST or 5% BSA in TBST at the indicated concentrations was performed, followed by three 10 min washing steps with TBST. Secondary antibodies diluted in 5% milk at indicated dilutions were then added onto the blot for 1 h, and then the blot was washed with TBST three times. When using HRP-conjugated secondary antibodies, Clarity Western ECL Substrate (Bio-Rad) was added to the blot for 5 min followed by imaging using the chemiluminescent channel of the ChemiDoc imaging system (Bio-Rad). Fluorescent and colourimetric channels of ChemiDoc (Bio-Rad) were used for imaging fluorescent secondary antibodies and protein ladders, respectively. To determine protein concentration, the BCA assay was used according to the protocol provided by ThermoFisher Scientific. Compatibility of the lysis buffer reagents determined which assay was used and absorbance readings were recorded at 562 nm, using the CLARIOstarPlus Microplate Reader (BMG LabTech).

### XL-BioID

The following steps, except for the BioLock treatment, were performed as described in Geoghegan et al. (2022). Briefly, *DUB2* and *CLK1* were endogenously tagged with the miniTurbo biotin ligase at their C- and N-termini, respectively, using CRISPR-Cas9 endogenous tagging (*60*). This was performed in the parental T7/Cas9 promastigote cell line. Double allele tagging was achieved using two different donor constructs containing either *BSD* or *PAC* resistance markers which were generated using the pPLOTv1 myc::miniTurboID::myc plasmids pGL2822 and pGL2835, respectively. The generated cell lines were called DUB2::miniTurbo::Myc and Myc::miniTurbo::CLK1. Primers are listed in **Supplementary Table 5.**

### BioLock treatment, biotin labelling and crosslinking

2 μl of BioLock Biotin blocking solution (IBA Lifescience) per mL of HOMEM was added in a log-phase promastigote cell culture for 16 h. Cells were pelleted (1,000 x *g*, 10 min, RT), washed with filtered PBS (Phosphate Buffered Saline), harvested (1,000 x *g*, 10 min, RT), and resuspended with fresh HOMEM at a cell density of 5 x 10^6^ cells mL^-1^.For *in vivo* biotin labelling, biotin solution (Merck) was added to a log-phase (BioLock-treated) promastigote cell culture at a final concentration of 500 μM for 30 min at 25°C. Samples were gently shaken every 5 min. Cells were then harvested (1,000 x *g*, 10 min, 4°C) and washed twice with ice-cold PBS (1,000 x *g*, 10 min, 4°C). *In vivo* cross-linking was then performed, by resuspending cells with 1 mM dithio(succinimidyl) propionate (DSP) crosslinker dissolved in pre-warmed (37°C) PBS and incubated for 10 min at 25°C. Cells were gently shaken every 2 min. To quench the cross-linking reaction, Tris-HCl pH 7.5 was added to a final concentration of 20 mM followed by incubation for 5 min at 25°C. Cells were harvested (1,000 x *g*, 10 min, RT) and stored at −80°C.

### Lysis, streptavidin pull-down and on-bead protein digestion

Frozen cells were thawed and resuspended in 500 μl of ice-cold radioimmunoprecipitation assay (RIPA) buffer supplemented with 0.1 mM PMSF, 1 μg mL^-1^ pepstatin A, 1 μΜ E-64 and 0.4 mM 1-10 phenanthroline. For every 3 mL of RIPA buffer, one cOmplete ULTRA Tablet (Mini, EASYpack Protease Inhibitor Cocktail, Roche) was added. Cells were then sonicated using the Bioruptor Pico sonication device (Diagenode) with 2 sonication cycles 30 s ON/30 s OFF, and samples were treated with 1 μl of Benzonase (250 Units, Abcam) for 1 h on ice. Cell extracts were then centrifuged (10,000 x *g*, 10 min, 4 °C), and the supernatant was retained. Biotinylated material was enriched by addition of magnetic streptavidin bead suspension (Resyn Bioscience), followed by end-over-end rotation o/n at 4°C. Beads were then washed successively for 5 min with an end-over-end rotation at room temperature with the following solutions: RIPA (4 x washes), 4 M urea in 50 mM triethyl ammonium bicarbonate (TEAB) pH 8.5 (1 x wash), 6 M urea in 50 mM TEAB pH8.5 (1 x wash), 1 M KCl (1 x wash) and 50 mM TEAB pH8.5 (1 x wash). On-bead digestion was then performed by re-suspending the beads in 50 mM TEAB pH 8.5 containing 10 mM TCEP, 10 mM iodoacetamide, 1 mM CaCl2, 1M urea and 500 ng trypsin/Lys-C (Promega) and incubation overnight at 37°C with shaking at 200 rpm. Supernatant containing trypsinised peptides was retained in LoBind protein eppendorf tubes. Beads were washed twice with deionised H2O for 5 min with end-over-end rotation at RT, and these were mixed with the retained supernatant. The resulting peptide mixtures were then acidified by addition of trifluoroacetic acid (TFA) to a final concentration of 0.5% v/v, centrifuged (17,000 x *g*, 10 min, RT), and the supernatant was kept for peptide desalting. Peptide desalting was performed, as described for enriched DiGly peptides.

### Liquid chromatography with tandem mass-spectrometry (LC-MS/MS)

Samples were resuspended in aqueous 0.1% v/v TFA solution and loaded onto an mClass nanoflow UPLC system (Waters) equipped with a nanoEaze M/Z Symmetry 100 Å C 18, 5 µm trap column (180 µm x 20 mm, Waters) and a PepMap, 2 µm, 100 Å, C 18 EasyNano nanocapillary column (75 mm x 500 mm, Thermo). The trap wash solvent was aqueous 0.05% (v:v) trifluoroacetic acid and the trapping flow rate was 15 µL/min. The trap was washed for 5 min before switching flow to the capillary column. Separation used gradient elution of two solvents: solvent A, aqueous 0.1% (v:v) formic acid; solvent B, acetonitrile containing 0.1% (v:v) formic acid. The flow rate for the capillary column was 300 nL/min and the column temperature was 40°C. The linear multi-step gradient profile was: 3-10% B over 8 mins, 10-35% B over 115 mins, 35-99% B over 30 mins and then proceeded to wash with 99% solvent B for 4 min. The column was returned to initial conditions and re-equilibrated for 15 min before subsequent injections.

The nanoLC system was interfaced with an Orbitrap Fusion Tribrid mass spectrometer (Thermo) with an EasyNano ionisation source (Thermo). Positive ESI-MS and MS2 spectra were acquired using Xcalibur software (version 4.0, Thermo). Instrument source settings were: ion spray voltage, 1,900 V; sweep gas, 0 Arb; ion transfer tube temperature; 275°C. MS 1 spectra were acquired in the Orbitrap with: 120,000 resolution, scan range: *m/z* 375-1,500; AGC target, 4e^5^; max fill time, 100 ms. Data dependent acquisition was performed in top speed mode using a 1 s cycle, selecting the most intense precursors with charge states >1. Easy-IC was used for internal calibration. Dynamic exclusion was performed for 50 s post precursor selection and a minimum threshold for fragmentation was set at 5e^3^. MS 2 spectra were acquired in the linear ion trap with: scan rate, turbo; quadrupole isolation, 1.6 *m/z*; activation type, HCD; activation energy: 32%; AGC target, 5e^3^; first mass, 110 *m/z*; max fill time, 100 ms. Acquisitions were arranged by Xcalibur to inject ions for all available parallelizable time.

### Data analysis of MS data

Peak lists in .raw format were imported into Progenesis QI (Version 2.2., Waters) to allow peak picking and chromatographic alignment. Concatenated MS2 peak list was exported in .mgf format and was searched through the Mascot programme (Matrix Science Ltd., version 2.7.0.1) against the TriTrypDB-derived *L. mexicana* subset database, which is composed of 8,250 sequences and 5,180,224 amino acid residues. The percolator algorithm was used for peptide identification to 1% FDR, and identified peptides were then imported into Progenesis QI, in which they were associated with precursor intensities. Non-conflicting, relative precursor intensities were normalized based on the total peptide signal per sample, and they were utilized for relative protein quantification. Quantifications were filtered to include only proteins with a minimum of two unique peptides. Next, data were transformed in Rstudio and proximal proteins were identified from the statistical package limma at 1% FDR.

### Alamar blue viability assay

Alamar Blue assays (Thermo Fisher Scientific) were performed to assess whether addition of a tag or BioLock treatment affected the metabolic activity of the parasites. 200 μl of promastigote cultures at a cell density of 1 x 10^6^ cells mL^-1^ were added in triplicate into wells of a 96-well plates. 125 μg mL^-1^ of resazurin dissolved in PBS was added to the wells and the plates were incubated for 8 h at 25°C. Metabolically active cells convert resazurin to resorufin. To detect this activity, fluorescence was recorded at 590 nm using the CLARIOstarPlus Microplate Reader (BMG LabTech). To calculate the percentage difference of fluorescence between sample and control cells, the following formula was used: (Average FI_590_ of treated sample / Average FI_590_ of untreated control)*100, of which FI_590_ is the intensity of fluorescence emission at 590 nm. Statistical analyses were performed in Rstudio.

For testing the extent to which tags affect parasite differentiation, axenic amastigote cells were prepared at a cell density of 1 x 10^6^ cells mL^-1^. 200 μl of axenic amastigote cell cultures were added in triplicate in 96-well plates, and incubated at 33°C. At 0 h, 24 h, 48 h and 72 h post-induction of promastigote-to-amastigote differentiation, 125 μg mL^-1^ of resazurin was added for 8 h at 33°C. Recording of fluorescence and data analysis was performed as in promastigote cells above.

### Co-immunoprecipitation

Co-immunoprecipitation (co-IP) assays were performed to test whether candidate DUB2 substrates form a stable protein complex with DUB2 and whether their protein abundance is affected by DUB2 depletion. DUB2 substrates were tagged endogenously with a 3xMyc epitope at either their N- or C-termini in *L. mexicana* DUB2^Flox+/+^ promastigotes. For co-IP, 30 mL of mid-log phase cultures were treated either with DMSO or 300nM rapamycin for 4 days, harvested, (1,200 x *g*, 10 min, RT), and resuspended in 10 mL of 1 mM DSP crosslinker dissolved in pre-warmed at 37°C PBS followed by incubation for 10 min at 25°C. To quench the cross-linking reaction, Tris pH 7.5 was added to a final concentration of 20 mM and the parasites were washed with PBS (1,200 x *g*, 10 min, RT). Cells were then resuspended in RIPA buffer supplemented with 250 units BaseMuncher Endonuclease and cOmplete ULTRA Tablet (Mini, EASYpack Protease Inhibitor Cocktail; 1 tablet per 3 mL of RIPA buffer; Roche). To ensure efficient lysis of the cells, sonication was also performed using the Bioruptor Pico sonication device (Diagenode) with 2 sonication cycles 30 s ON/30 s OFF. Cell extracts were then centrifuged (10,000 x *g*, 10 min, 4°C), and 20 μl of the supernatant was extracted and prepared for Western Blot analysis as a pre-immunoprecipitation sample (whole cell lysate; WCL). The rest of the supernatant was immunoprecipitated using 30 μl anti-Myc magnetic beads (Pierce) for 2 h with end-over-end rotation at 4°C. The anti-Myc magnetic beads were washed twice with a PBS buffer supplemented with 50 mM NaH_2_PO_4_ pH 7.5, 150 mM NaCl and 0.025% Tween. Beads were then washed three times in lysis buffer for 5 min with end-over-end rotation at room temperature. Proteins were then eluted using 1 x LDS buffer supplemented 250 mM DTT followed by a 10 min incubation at 70°C. The presence of bait and prey proteins was investigated by Western Blotting and probing for the Myc and HA epitopes, respectively.

### Harvesting, lysis and protein digestion for ubiquitinomics

Promastigotes and amastigotes were grown to cell densities of 4 x 10^6^ cells mL^-1^ and 1 x 10^7^ cells mL^-1^ before 1.5 x 10^9^ cells and 4 x 10^9^ cells were pelleted (1,200 x *g*, 10 min, RT), respectively. Following two washes with PBS, cells were lysed in 5% w/v SDS buffer (SDS; Thermo Fisher Scientific, 50 mM TEAB buffer; Merck) and sonicated with a microtip sonicator (Sonics Vibra-Cell) in two 10 s pulses at an amplitude of 50% separated by a 1 min pause on ice. Due to the high viscosity of the lysate, additional sonication was performed using the Bioruptor Pico sonication device (Diagenode) with 2 sonication cycles 30 s ON/ 30 s OFF, at 4°C. The lysate was centrifuged for 15 min at 10,000 x *g* and the supernatant was transferred to a fresh protein LoBind eppendorf tube. Protein quantification was performed using the Pierce BCA assay (23225; Thermo Fisher Scientific), following which equal amounts of total protein were used in each of the downstream experiments. To reduce protein disulphide bonds, dithiothreitol (DTT) was added to a final concentration of 1 mg mL^-1^ with incubation at 55°C for 1 h with stirring at 350 rpm. The lysate was then cooled to room temperature on ice, and proteins were alkylated by addition of a 1/10 volume of 100 mM 2-chloroacetic acid (Merck) and incubation at room temperature in the dark for 30 min. At this point, 10 μg of protein sample was removed (pre-trypsinisation sample), mixed with 1 x LDS buffer (NuPAGE; NP0007; Thermo Fisher Scientific) containing 10 mM DTT (R0861; Thermo Fisher Scientific), and incubated at 70°C for 10 min before storage at −20°C.

To the remaining lysate, a 1/10 volume of 12% phosphoric acid (Merck) was added and the solution was mixed thoroughly. Next, 6.6 volumes of 90% methanol (Honeywell)/100 mM TEAB (Merck) buffer was added to the lysate, and a cloudy protein colloid was formed. Immediately, the sample was transferred onto an S-Trap Midi Cartridge (≥ 300 μg; ProtiFi). The cartridge was centrifuged for 2 min at 4,000 x *g*, at room temperature and the flow-through was discarded. To desalt the sample, the S-Trap cartridge was washed 4 times with 90% methanol/100 mM TEAB buffer and centrifuged as before. The cartridge was next transferred into a 15 mL falcon tube. For protein digestion, trypsin/Lys-C protease mix (Mass Spec Grade; V5071; Promega) was resuspended in 50 mM TAEB (pH ∼8) at a concentration of 1:50 protease mix: protein sample was added onto the S-Trap cartridge, which was incubated overnight at 37°C. The falcon tube was placed vertically to ensure even absorption of the protease mix and the lid of the tube was loosely closed to allow absorption onto the cartridge without creating negative pressure. The following day, three steps of centrifugation for 2 min at 4,000 x *g* at room temperature were carried out following addition of firstly 50 mM TAEB buffer, secondly 0.5% v/v TFA in water (Merck) and thirdly 50% v/v ACN (Thermo Fisher Scientific)/ 0.5% v/v TFA in water. The flow through solutions were retained. The pH of the eluate was measured and it was > 3.5, 50% v/v TFA in water was added. 10 μg of peptide mixture was removed and treated as a pre-trypsinisation sample and used later in stain-free gel imaging to assess the success of the protease digestion. 100 μg of peptides from each sample were dried overnight in a vacuum concentrator (Speed-Vac) at 37°C and stored at −80°C. The former samples were used for total proteomics, while the latter samples were further processed for total ubiquitinomics. All of the MS-related experiments were conducted using filter tips, Protein LoBind tubes (Eppendorf) and MS-grade water (Thermo Fisher Scientific).

### K- ε -GG remnant immunoaffinity purification

Immunoaffinity purification was conducted according to the PTMScan HS K-ε -GG remnant magnetic immunoaffinity beads protocol (Cell Signalling Technology). Briefly, the lyophilized peptides were centrifuged for 5 min at 2,000 x *g*, at room temperature and resuspended in 1 X HS IAP Bind buffer. The pH of the dissolved peptides was adjusted to pH 7 by addition of 1M Tris base as required. The tube was gently shaken for 5 min, at room temperature (<500 rpm). The solution was cleared by centrifugation for 5 min at 10,000 x *g* at 4°C and further cooled on ice. Meanwhile, the antibody-conjugated magnetic beads were washed and prepared before adding the samples, as follows. The vial of antibody-bead slurry was centrifuged for 2 sec at 2,000 x *g*, and the beads gently resuspended by pipetting. The bead slurry was placed into a protein LoBind eppendorf tube. Ice-cold PBS was added to the beads with mixing by inverting the tube five times. The tube was then placed on a magnetic rack for 10 sec and the buffer was carefully removed. This washing step was repeated four times. The soluble peptide fractions were transferred into the tube containing the antibody beads and incubated for 2 h on an end-over-end rotator, at 4°C. Sequentially, the tube was centrifuged for 2 sec at 2,000 x *g*, placed on a magnetic rack for 10 sec and the unbound peptide solution was discarded. Beads were resuspended in HS IAP Wash buffer, mixed by inverting the tube five times, centrifuged for 2 sec at 2,000 x *g*, and the buffer was removed upon placing the tube for 10 sec on a magnetic rack. This washing step was repeated for another three times followed by an additional two washes with chilled LCMS water. To elute the bound peptides from the antibodies bound beads, 0.15% v/v aqueous TFA was added with gentle mixing at room temperature. The tube was centrifuged at 2,000 x *g* for 2 sec and the eluted sample was transferred to a fresh protein LoBind eppendorf tube. This step was repeated once more prior to desalting.

### Peptide desalting

C18 desalting tips were placed into a protein LoBind eppendorf tube, and equilibrated successively with 100 μl of the following solutions: aqueous 100% v/v acetonitrile (ACN) solution, aqueous 80% v/v ACN 0.1% v/v TFA solution, and aqueous 0.1% v/v TFA solution. Upon adding each washing solution, the C18 tips were spun down, and then the flow through was discarded. Peptide-containing supernatants were loaded onto the equilibrated C18 tips with the flow-through recirculated three times onto the column. The C18 tips were washed twice with 0.1% v/v TFA solution, and peptides were eluted with 2 x 30 μl of aqueous 80% v/v ACN 0.1% v/v TFA solution. The combined eluates were mixed and lyophilised for MS analysis. All the centrifugation steps in this section were performed at 2,000 rpm for 1-2 min at RT.

### timsTOF MS data acquisition and data analysis

The following approach was taken for both total proteomics- and ubiquitinomics-based timsTOF mass spectrometry. Lyophilised peptides were resuspended in aqueous 0.1% TFA solution and loaded onto EvoTip Pure tips for desalting and as a disposable trap column for nanoUPLC using an EvoSep One system. A pre-set EvoSep 15 SPD gradient (Evosep One HyStar Driver 2.3.57.0) was used with a 15 cm EvoSep C_18_ Performance column (15 cm x 150 mm x 1.5 mm).

The nanoUPLC system was interfaced to a timsTOF HT mass spectrometer (Bruker) with a CaptiveSpray ionisation source (Source). Positive PASEF-DDA, ESI-MS and MS^2^ spectra were acquired using Compass HyStar software (version 6.2, Bruker). Instrument source settings were: capillary voltage, 1,600 V; dry gas, 3 l/min; dry temperature; 180°C. Spectra were acquired between *m/z* 100-1,700. TIMS settings were: 1/K0 0.6-1.60 V.s/cm^2^; Ramp time, 100 ms; Ramp rate 9.42 Hz. Data dependent acquisition was performed with 10 PASEF ramps and a total cycle time of 1.9 s. An intensity threshold of 2,500 and a target intensity of 20,000 were set with active exclusion applied for 0.4 min post precursor selection. Collision energy was interpolated between 20 eV at 0.6 V.s/cm^2^ to 59 eV at 1.6 V.s/cm^2^.

Data were searched using FragPipe (v22) (*61*) against the *L. mexicana T7/Cas9* derived (*62*) proteome database appended with common proteomic contaminants. Search parameters specified trypsin protease with up to three missed cleavage positions. Mass tolerance was set to 15 ppm for MS1 and MS2 matching. For variable modifications, oxidised of methionine, protein N-terminal acetylation and GlyGly (GG) derivatisation of lysine (+114.04293 Da) were selected. Validation was performed using ProteinProphet and PeptideProphet within Philosopher (v5.1). Searches were run at 1% FDR for both peptide and protein as assessed empirically against a reverse database search. Site localisation probabilities were calculated with PeptideProphet. Intensities from accepted peptides were extracted using IonQuant (v1.10).

For the ubiquitinomics data, ubiquitination sites with a localization possibility less than 0.5 were excluded, and for the proteomics datasets, only proteins with at least two unique peptides were kept for downstream analysis. Further protein/peptide filtering, data processing and statistical analysis to identify significantly enriched ubiquitination sites or proteins was performed using limma via FragPipe-Analyst (*61*). Specifically, the following parameters were set at the indicated values: 0.75 for the minimum percentage of non-missing values in at least one condition, 0.05 for the differentially expressed adjusted *p*-value cutoff, and 1 for the differentially expressed Log2 fold change cutoff. Variance stabilizing normalization was selected as normalization type, and for imputation and type of FDR correction min and Benjamini Hochberg were applied, respectively.

## Acknowledgments

This work was supported by funding from the Wellcome Trust (223045/Z/21/Z). The York Centre of Excellence in Mass Spectrometry was created thanks to a major capital investment through Science City York, supported by Yorkshire Forward with funds from the Northern Way Initiative, and subsequent support from EPSRC (EP/K039660/1; EP/M028127/1). We thank our colleagues in The Bioscience Technology Facility of the University of York, who provided expertise and technical support, and Andreas Damianou (University of Oxford) for his critical input related to the *Leishmania* DUBs and the design of the ubiquitinomics experiments.

## Disclosure and competing interests statement

The authors declare that they have no conflict of interest

## Data availability

Mass spectrometry data sets and proteomic identifications are available to download from MassIVE (MSV000101252) and are referenced in ProteomeXchange (PXD076194).

## Author contributions

J.C.M., C.M., and A.J.W. acquired funding and supervised the project.

S.A., J.C.M., C.M., and A.J.W conceived the project.

S.A., A.J.W and J.C.M. designed the experiments.

S.A, A.D., R.N., and C.Mc. performed the experiments.

S.A., A.D., V.G., and N.G.J. analysed experimental data.

S.A., N.G.J and JCM drafted the manuscript and all other authors reviewed it.

